# Fatherhood alters gene expression within the MPOA

**DOI:** 10.1101/258111

**Authors:** Adele M. H. Seelke, Jessica M. Bond, Trent C. Simmons, Nikhil Joshi, Matthew L. Settles, Danielle Stolzenberg, Mijke Rhemtulla, Karen L. Bales

## Abstract

Female parenting is obligate in mammals, but fathering behavior among mammals is rare. Only 3–5% of mammalian species exhibit biparental care, including humans, and mechanisms of fathering behavior remain sparsely studied. However, in species where it does exist, paternal care is often crucial to the survivorship of offspring. The present study is the first to identify new gene targets linked to the experience of fathering behavior in a biparental species using RNA sequencing. In order to determine the pattern of gene expression within the medial preoptic area that is specifically associated with fathering behavior, we identified differentially expressed genes in male prairie voles (*Microtus ochrogaster)* that experienced one of three social conditions: virgin males, pair bonded males, and males with fathering experience. Differentially expressed genes from each comparison (i.e., Virgin vs Paired, Virgin vs Fathers, and Paired vs Fathers) were evaluated using the Gene Ontology enrichment analysis, and Kegg pathways analysis to reveal metabolic pathways associated with specific differentially expressed genes. Using these tools, we identified a group of genes that are differentially expressed in voles with different amounts of social experience. These genes are involved in a variety of processes, with particular enrichment in genes associated with immune function, metabolism, synaptic plasticity, and the remodeling of dendritic spines. The identification of these genes and processes will lead to novel insights into the biological basis of fathering behavior.

---

**Declarations of Interest:** none

## 1. Introduction

Biparental care, where both mother and father contribute to the care of the offspring, is displayed by a minority of mammalian species – usually cited as 3–5% (Kleiman, 1977; Lukas and Clutton-Brock, 2013; Opie et al., 2013). Female parenting is obligate because mammalian offspring need to nurse. Therefore, the presence of the male, and particularly active participation of the male in fathering, is an unusual situation seen only in our own and a limited number of other mammalian species (Gubernick and Alberts, 1987; Mendoza and Mason, 1997; Runcie, 2000; Thomas and Birney, 1979; Wynne-Edwards and Timonin, 2007). In species where it does exist, including humans, paternal care is often crucial to the survivorship of offspring, or at the least has significant and long-term impacts on growth as well as neural, reproductive and social development (Bales and Saltzman, 2016; Cantoni and Brown, 1997); however little is known about the specific neurobiological regulation of paternal care (Wynne-Edwards and Timonin, 2007). The vast majority of parenting research focuses on the mother, while the role of the father is mostly considered in the context of paternal absence (Bales and Saltzman, 2016). Considering paternal care through the absence of the father in a biparental species has drawbacks, however, since it is impossible to distinguish between the influence of the quantitative absence of another caregiving individual and the qualitative absence of the father in particular.

However, paternal absence is the most extreme situation. Although less studied, individual variation in fathering can also have long-term effects on offspring (Bales and Saltzman, 2016), and in the context of non-human mammals is always carried out in a biparental care situation. In prairie voles, we have shown that natural variation in biparental parenting behavior predicts pup development and juvenile social behavior (Perkeybile et al., 2013), exploratory behavior and pair-bonding, and adult aggression and stress responses (Arias Del Razo and Bales, 2016; Perkeybile and Bales, 2015a; Perkeybile and Bales, 2015b). It is not always possible in a biparental care situation to tell what outcomes in offspring are due to maternal care and what are due to paternal care. However, some very interesting roles for the father have been observed. For instance, in some species, males may compensate for poor maternal care (or allow mothers to expend less energy on non-nutritive tasks like carrying)(Bales et al., 2002; Perkeybile et al., 2013); or a paternal behavior such as retrievals (carrying pups back to the nest or territory) may be directly linked to offspring display of retrievals and aggression as an adult (Bester-Meredith and Marler, 2003; Frazier et al., 2007).

While little is known about the effects of paternal care on offspring, especially when compared to maternal care, even less is known about the neural mechanisms underlying fathering behavior. It has been hypothesized that similar neural circuits are responsible for both maternal and paternal behaviors (Dulac et al., 2014). While alterations in neural activity appear to be hormonally regulated in females, hormonal manipulation in males does not have the same effect on paternal behavior (Saltzman and Ziegler, 2014). This has led some to suggest that maternal and paternal behavior depends upon non-homologous neuroendocrine circuits (Wynne-Edwards and Timonin, 2007), and has raised the question of what factors are involved in the generation of these behaviors.

Although the neuroendocrine contributions to parenting may vary by sex, it is believed that the neural circuit governing parental behavior is conserved across sex (Kohl et al., 2018). The MPOA is a central node in the neural circuit that regulates both maternal and paternal care and has long been recognized as playing a critical role in the generation and regulation of parental behavior (see (Bales and Saltzman, 2016; Kohl and Dulac, 2018) for review). Paternal experience increases Fos immunoreactivity in the MPOA of California mice (de Jong et al., 2009). Virgin male prairie voles that were exposed to pups also showed an increase in Fos immunoreactivity within the MPOA (Kirkpatrick et al., 1994). Lesions of the MPOA disrupt both maternal and paternal behavior in biparental California mice (Lee and Brown, 2002), and maternal behavior in rats (Numan et al., 1988) and mice (Tsuneoka et al., 2013).

The relationship between maternal behavior and gene expression has been examined in a variety of species. The initiation of maternal behavior is reduced in oxytocin knock-out mice (Rich et al., 2014). In humans, differences in maternal responsiveness to babies and toddlers were associated with variants in serotonin transporter and oxytocin receptor genes (Bakermans-Kranenburg and van Ijzendoorn, 2008; Feldman et al., 2012). Maternal experience in mice was linked to large scale changes in gene expression in the lateral septum (Eisinger et al., 2013). And mice that were bred for high or low levels of maternal aggression showed a wide range of differentially expressed genes within the hypothalamus (Gammie et al., 2007). It should be noted that these studies only examined mothers and the majority targeted a specific gene or genes that were already implicated in parental behavior or attachment. As such, they did not identify or examine novel gene targets that may also play a role in fathering behavior.

The goal of this study was to identify novel gene targets and potential mechanisms that may contribute to the production and regulation of paternal behavior. We analyzed gene expression in three groups of adult male prairie voles: virgin males, males who had formed a pair bond with a female, and males who had fathering experience. Samples were taken from the medial preoptic area (MPOA), a region that is central to the expression of both maternal and paternal behaviors (Dulac et al., 2014; Kuroda and Numan, 2014; Rilling and Young, 2014; Stolzenberg and Champagne, 2016), and RNA was extracted and sequenced.

## 2. Materials and Methods

### 2.1 Subjects

Subjects were 18 adult male prairie voles. Animals were born and housed in the Psychology Department Vivarium at the University of California, Davis. These animals were descendants of a wild stock originally caught near Champaign, Illinois. The animals were weaned at 20 days of age and pair housed with an animal of the same sex (sibling if available, similarly aged non-sibling if not) in small laboratory cages (27 × 16 × 13 cm) in which food and water were available *ad libitum*. All animals were maintained at approximately 70°F (21°C) on a 14:10 light/dark cycle with the lights on at 6 a.m. All experiments were performed under National Institutes of Health guidelines for the care of animals in research and were approved by the Institutional Animal Care and Use Committee of the University of California, Davis.

At postnatal day (P) 42–45 subjects were placed in one of three groups of age-matched males: 1) virgin males, 2) sexually experienced, “pair-bonded” males, or 3) males with fathering experience (Figure 1). This was designed to dissociate alterations in gene expression that were related to pair bonding from alterations related to paternal behavior. Virgin males were housed with a male same-age conspecific, and they were euthanized without engaging in sexual contact with females. Pair-bonded males were housed with a same-age female conspecific for ~20 days, after which the males were euthanized. Because mating and pregnancy strengthens pair bonds in prairie voles (Insel et al., 1995; Williams et al., 1992; Winslow et al., 1993; Young and Wang, 2004), we confirmed that females were pregnant. Pair-bonded males were euthanized before females gave birth, ensuring they had no contact with pups. The third group consisted of males which had ~3 days of paternal experience. These males were also housed with female pair-mates with whom they presumably formed a pair-bond. The females gave birth, and the males were permitted three days of contact with pups before they were euthanized. Three days of parental experience was chosen to minimize age differences between subjects. Furthermore, prairie vole fathers already exhibit large amount of paternal care by postnatal day 3 (Oliveras and Novak, 1986; Perkeybile et al., 2013).

**Figure 1:**
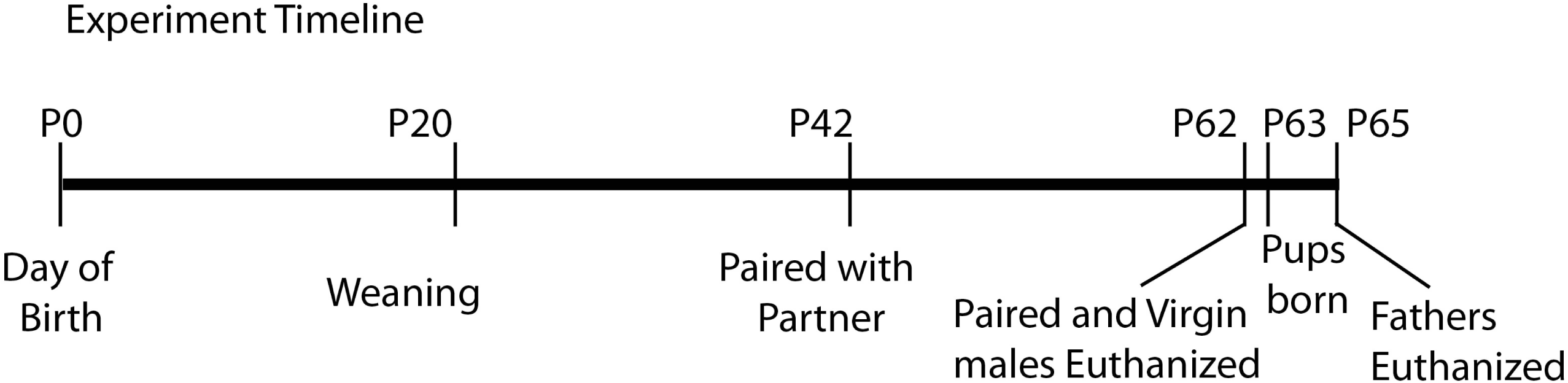
Timeline of social interactions across the lifespan. All subjects were separated from their parents and removed from their home cage at postnatal day (P) 20 and housed with a same-sex conspecific. Around P42, males assigned to the “paired” and “father” groups were rehoused with an opposite-sex conspecific, while males in the “virgin” group remained with their same-sex conspecific. 20 days later (~P62), the subjects in the “virgin” and “paired” groups were euthanized and their brains were removed. One day later (~P63), pups were born to the males in the “father” group, and these males were euthanized 2–3 days later (~P65).

Subjects were anesthetized using isoflurane and euthanized via cervical dislocation. Upon euthanasia, brains were removed and flash frozen. The brains were sliced on a cryostat into 120 µm sections and mounted on slides. Punches were taken from the MPOA using a 15.5-gauge blunt needle (Figure 2), and were stored in a −80 freezer until RNA extraction. The sequence of analyses for this study is outlined in Figure 3.

**Figure 2:**
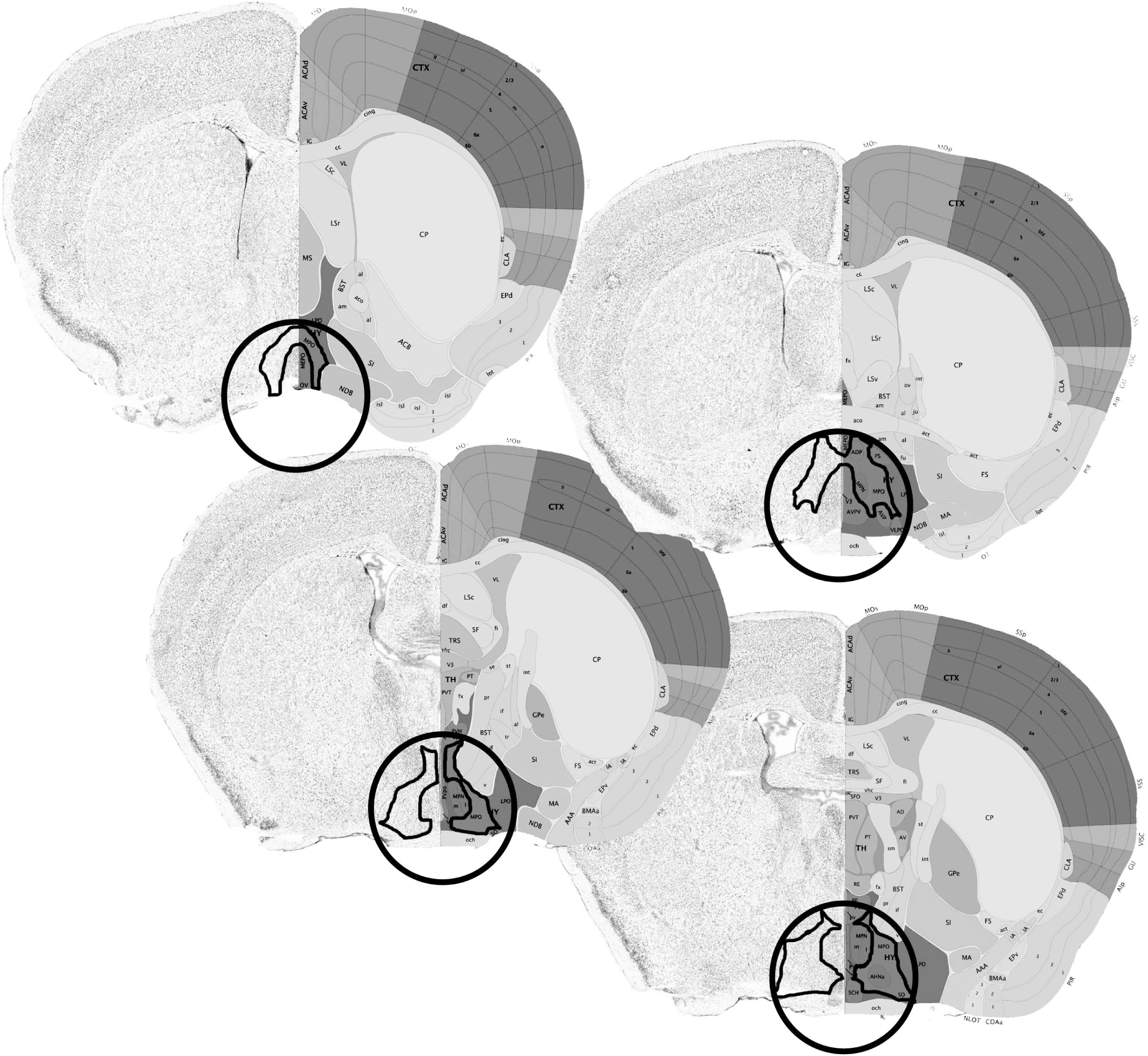
A schematic representing the area from which tissue samples were taken. The circumference of the tissue punch is delineated by a circle, and the MPOA is outlined in black. The tissue punches removed the entirety of the MPOA, as well as small portions of adjacent hypothalamic tissue.

**Figure 3:**
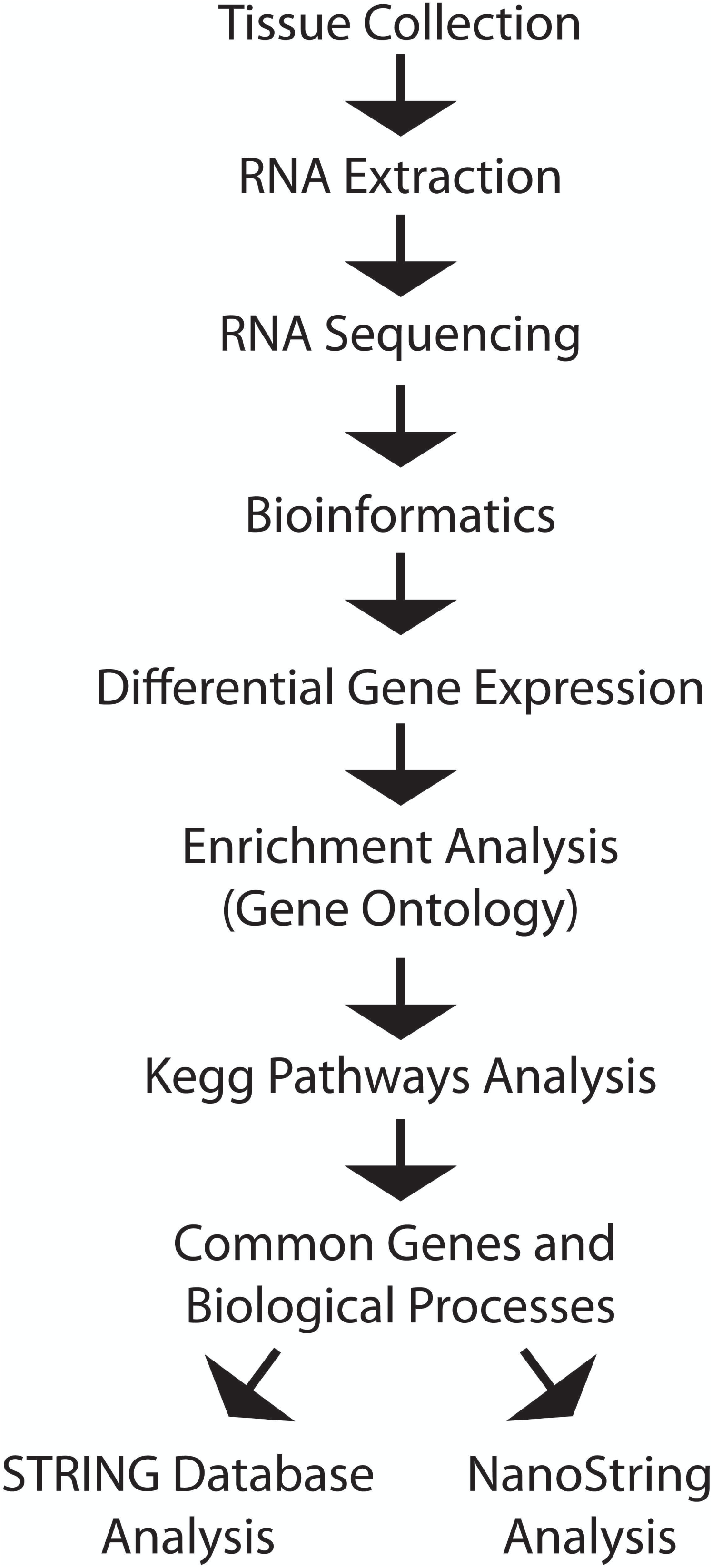
Analysis plan for RNA sequencing experiment. Following the collection of brain tissue, RNA was extracted from the MPOA and sequenced at the UC Davis Expression Analysis Core facility. The resulting sequences were linked to specific genes and analyzed for relative frequency at the UC Davis Bioinformatics Core facility. The top 500 differentially expressed genes in each comparison group were analyzed using the Gene Ontology database (geneontology.org) and Kegg Pathways analysis (www.kegg.jp). These analyses allowed us to identify a subset of genes of interest and their associated biological processes that were differentially expressed in each comparison group. We then analyzed the interrelatedness of these groups of genes using STRING Database analysis (string-db.org). Finally, the expression of 30 genes of interest was quantitatively expressed using the NanoString Analysis.

### 2.2 RNA Extraction

Total RNA was isolated with Qiazol reagent (Qiagen) and purified with an RNeasy^®^ Plus Micro Kit (74004; Qiagen, Valencia, CA) as well as the optional DNase digestion (Qiagen 129046). A Nanodrop™ Spectrophotometer was used to determine the quality and quantity of the RNA. All samples had a 260/280 ratio > 1.8.

### 2.3 RNA Sequencing

A total of 18 RNA-seq libraries were prepared from the RNA of the 18 male prairie voles (Table 1). RNA sequencing and library preparation was performed by the DNA Technologies and Expression Analysis Core in the Genome Center of the University of California, Davis. Barcoded RNA-seq libraries were generated from 1 ug total RNA each after poly-A enrichment using the Kapa Stranded RNA-seq kit (Kapa Biosystems, Cape Town, South Africa) following the instructions of the manufacturer. The libraries were generated on a Sciclone G3 liquid handler (Caliper Life Sciences, Alameda, CA). Quality was verified with the Bioanalyzer 2100 instrument (Agilent, Santa Clara, CA) and quantified by fluorometry on a Qubit instrument (LifeTechnologies, Carlsbad, CA) and pooled in equimolar ratios. The pooled library was then quantified by qPCR with a Kapa Library Quant kit (Kapa Biosystems) and sequenced on 1 lane of an Illumina HiSeq 4000 (Illumina, San Diego, CA) with paired-end 150bp reads.

**Table 1:** RNA data

| ID | Raw Reads | Trimmed Reads | % Reads Kept | % Aligned | % Aligned to rRNA |
| --- | --- | --- | --- | --- | --- |
| V1 | 25764898 | 25657648 | 99.58373598 | 95.1 | 5.67 |
| V2 | 20747099 | 20665472 | 99.60656186 | 95.99 | 2.63 |
| V3 | 23633483 | 23500072 | 99.43550005 | 96.13 | 2.07 |
| V4 | 17827384 | 17709907 | 99.34103063 | 96.06 | 2.38 |
| V5 | 22459377 | 22360940 | 99.56171091 | 95.59 | 4.09 |
| V6 | 22690769 | 22591612 | 99.56300732 | 96.09 | 2.78 |
| P1 | 22451193 | 22317681 | 99.40532336 | 92.7 | 8.72 |
| P2 | 23931482 | 23837973 | 99.60926365 | 95.84 | 2.26 |
| P3 | 21258757 | 21155922 | 99.51626993 | 94.97 | 5.00 |
| P4 | 20114070 | 20011415 | 99.48963586 | 95.86 | 3.16 |
| P5 | 21770575 | 21677246 | 99.57130668 | 96.11 | 2.06 |
| P6 | 19282593 | 19185302 | 99.49544649 | 95.73 | 3.28 |
| F1 | 20833672 | 20702987 | 99.3727222 | 95.14 | 4.82 |
| F2 | 21680207 | 21568888 | 99.48654088 | 95.99 | 2.53 |
| F3 | 20565896 | 20479195 | 99.57842342 | 96.06 | 1.65 |
| F4 | 19759347 | 19680046 | 99.59866589 | 96.17 | 2.27 |
| F5 | 21472445 | 21367346 | 99.51054014 | 95.95 | 3.06 |
| F6 | 22055401 | 21961628 | 99.57482977 | 96.31 | 2.32 |

Raw sequencing data have been deposited at NCBI’s Sequence Read Archive (SRA) under study accession number SRP128134.

### 2.4 Bioinformatic analysis

Bioinformatic analysis was performed by the UC Davis Bioinformatics Core Facility also in the Genome Center. Briefly, reads were trimmed for adapter contamination and quality using scythe (version c128b19) and sickle (version 7667f147e6) respectively. The reads were then aligned to the prairie vole genome (MicOch1.0) using bwa mem (version 0.7.13), after which featureCounts (version 1.5.0-p1) was used to create the raw gene expressions counts. Finally, R (version 3.3.2) with the edgeR and limma/voom packages were used to filter and transform (voom transformation), and test for statistical significances between groups. Briefly, data were prepared by first choosing to keep genes that achieved at least 0.5 count per million (cpm) in at least five samples, normalization factors were calculated using trimmed mean of M-value (TMM), and the voom transformation was applied. A completely randomized design was implemented, comparisons of interest were extracted using contrasts, and moderated statistics were computed using the empirical bayes procedure eBayes. Finally, each gene was corrected for multiple testing using the Benjamini-Hochberg (BH) false discovery rate correction.

### 2.5 Gene Ontology Analysis

Differential gene expression was directly compared between each pair of groups, resulting in three comparisons: Virgin males vs Paired males (V vs P), Virgin males vs Fathers (V vs F), and Paired males vs Fathers (P vs F). None of the differentially expressed genes reached the level of statistical significance. In order to capture the genes that were most likely to show functional differentiation between comparison groups we performed gene ontology annotation enrichment analysis. The gene enrichment analysis annotated the differentially expressed genes using one of three data sets: cellular component, molecular function, or biological process. Enrichment testing was conducted using Kolmogorov-Smirnov testing as implemented in the Bioconductor package topGO (Alexa and Rahnenfuhrer, 2016). We next examined the GO annotations that were significantly enriched (raw P value < 0.05) and selected the GO annotations in each comparison that were related to the brain or behavior, excluding unrelated annotations (i.e. GO:0003014, Renal system process or GO:0008354, Germ cell migration). We then categorized the remaining annotations based on gross function within each comparison group.

### 2.6 Kegg Pathways Analysis

We examined the functions with the highest fold enrichment (# observed genes/# expected genes, fold enrichment >|1|, p < 0.05), then identified individual genes associated with that function. We ran each individual gene through the Kegg Pathways database (Kyoto Encyclopedia of Genes and Genomes; http://www.genome.jp/kegg/) to identify molecular signaling pathways associated with that gene. A single gene may be involved in a number of different metabolic and biological pathways, so we then identified commonly recurring pathways associated with the individual differentially expressed genes. Pathways that were unrelated to brain function (for example, those that were involved in kidney, liver, or heart metabolism) were not included in the analysis. When a specific gene was associated with multiple pathways of interest it was identified as a candidate gene for further analysis. For example, *Grin2a* was associated with 6 pathways that are involved in neural plasticity.

Using the qualitative procedures described previously, we identified 49 candidate genes which display differential expression. We then averaged gene expression across animals within each condition and transformed the data into ratios; the values we used for all analyses were the ratios of gene expression in each condition relative to the virgin condition. The expression ratio of genes in virgin animals was set at 1, a value >1 indicated that genes were more expressed relative to virgins, and a value <1 indicated that genes were less expressed relative to virgins. Thus, we analyzed whether gene expression for each gene varied between paired animals and fathers. Effect size was measured using Cohen’s *d*.

### 2.7 Assessment of gene interaction networks

After identifying each set of differentially expressed genes, we analyzed the connectivity of the gene network using the STRING Database (string-db.org) (Szklarczyk et al., 2017). The STRING database identifies protein-protein interactions between members of a gene set, which allows the user to build a network of functional gene interactions. STRING also measures the functional and interaction enrichments of the gene network, calling upon GO Annotations, Kegg pathways, and connections between nodes.

### 2.8 NanoString Analysis

Following the identification of differentially expressed candidate genes, we performed a quantitative analysis of the expression of 33 genes (30 target genes and 3 housekeeping genes) using the nCounter SPRINT profiler (NanoString Technologies, Seattle, WA). Genes were chosen to be included in the NanoString analysis based on their differential expression values as determined by the raw expression data, as well as their functional significance. One additional gene, *Bdnf*, was chosen due to previous studies indicating that it plays a significant role in plasticity and parenting (i.e., *Bdnf*) (Pereira, 2016; Tabbaa et al., 2017). The nCounter analysis assay was conducted using RNA that remained after the completion of the sequencing experiment.

Briefly, NanoString is a medium-throughput method that can analyze many genes within a single sample with comparable sensitivity and accuracy to quantitative real-time RT-PCR (Geiss et al., 2008). NanoString designed and manufactured custom probes corresponding to 33 genes we identified for quantitative analysis, consisting of 30 target genes and 3 housekeeping genes (*Gusb*, *Pgk1*, and *Eif4a2*). A codeset specific to a 100-base region of the target mRNA was designed using a 3’ biotinylated capture probe and a 5’ reporter probe tagged with a specific fluorescent barcode. Data were collected using the nCounter Digital Analyzer by counting the number of individual barcodes.

Each transcript of interest was recognized by a capture probe and a reporter probe, each containing 30–50 bases complementary to the target mRNA. To minimize assay variability, the code sets also included negative and positive control reporter probes that were developed by the External RNA Control Consortium (ERCC). Six positive control reporter probes (ERCC-selected mRNA targets) were pre-mixed with (Spike-Ins) the code set at a concentration range (0.125–128 fM), a range corresponding to the expression levels of most mRNA of interest, to control for overall efficiency of probe hybridization and determine the detection range for transcripts of interest in each assay. A scaling factor was calculated for each sample, and a scaling factor outside the range of 0.3 to 3 indicated suboptimal hybridization. In our samples, the scaling factor always fell within the optimal range and was thus applied to all counts in the sample.

Quantitative expression data from the nCounter was downloaded and analyzed using the nSolver software package (NanoString Technologies, Seattle, WA). The raw counts for all transcripts were multiplied by the scaling factor to produce the adjusted counts. The relative expression was determined for each comparison group, and the effect size of the difference between expression values was determined using Cohen’s *d*. Differential expression was also compared using t-tests, and p-values were adjusted for multiple comparisons in nSolver.

## 3. Results

### 3.1 Gene Ontology Analysis

Individual genes are associated with gene ontology annotations in order to describe the various functions of a particular gene product. The cellular component analysis describes the locations of gene expression, at the levels of subcellular structures. The molecular function analysis describes the function that each gene product performs within the cell. The Biological Process Analysis describes a recognized series of events or collection of molecular functions associated with a gene or gene product. Each analysis was completed for all differentially expressed genes in each of the three comparison groups, Virgin vs Paired (*V vs P*), Virgin vs Father (*V vs F*), and Paired vs Father (*P vs F*). Because each GO Annotation references many genes, in some instances the same GO Annotation was present in multiple comparison groups.

The initial GO enrichment analysis returned 209 GO Annotations in the *V vs P* comparison, 222 annotations in the *V vs F* comparison, and 264 in the *P vs F* comparison that were significantly enriched. Upon selecting the GO annotations in each comparison that were related to the brain or behavior, we were left with 47 GO Annotations in the *V vs P* comparison, 47 annotations in the *V vs F* comparison, and 61 annotations in the *P vs F* comparison. (Tables 2–4). We then categorized these annotations based on gross function (Figure 4). The functional categories of GO annotations were differentially distributed across the three comparison groups. Annotations related to Neuropeptide activity were only found in the *V vs P* comparison, whereas Immune Function annotations were most predominant in the *V vs F* comparison. The P vs F comparison contained the greatest number of annotations related to Plasticity, DNA/RNA/Transcription, and Axon/Dendrite/Synapse.

**Figure 4:**
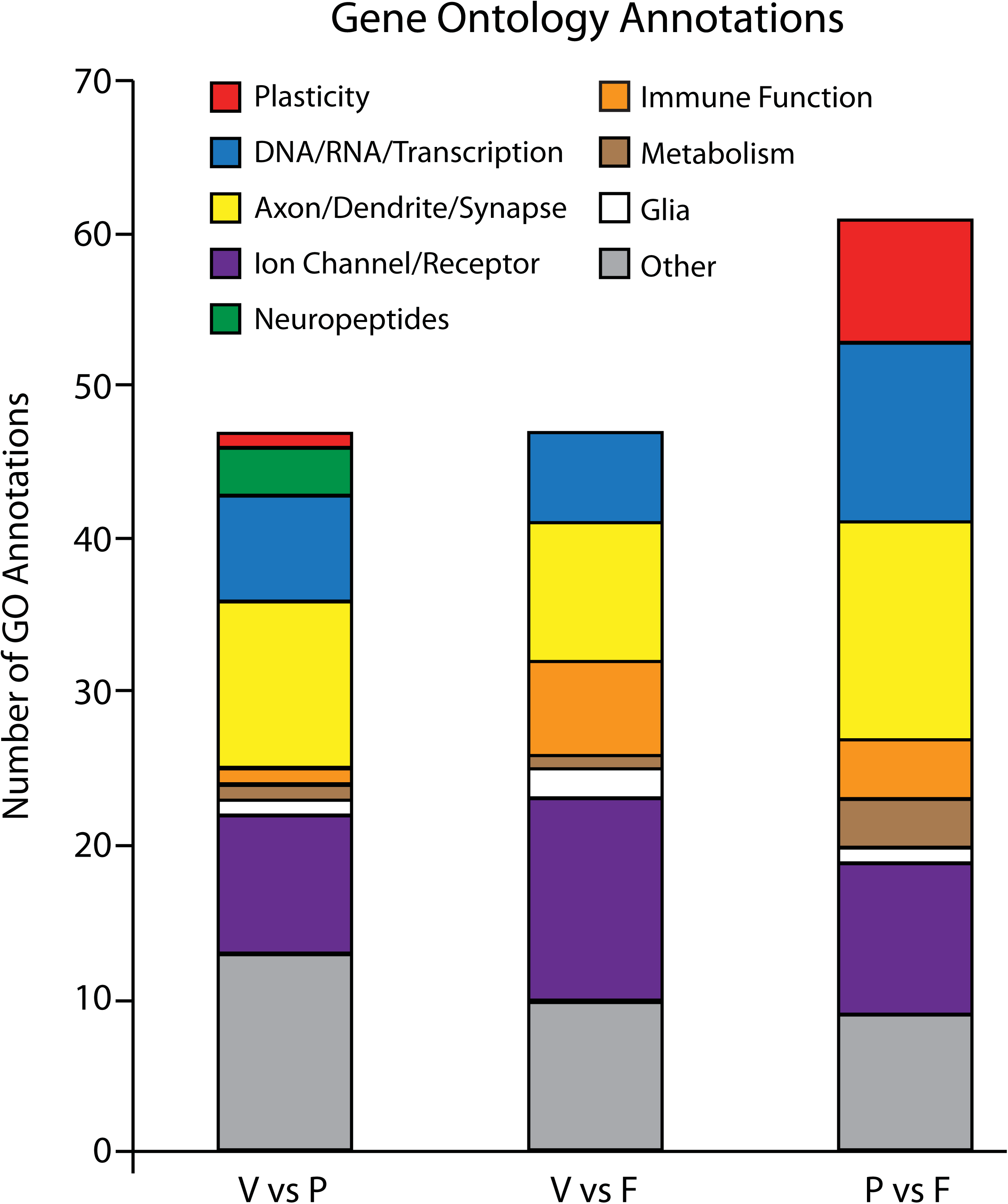
Enrichment of Gene Ontology Annotations across comparison groups. The gene enrichment analysis grouped the differentially expressed genes using gene ontology annotations data. We selected significantly enriched GO annotations and identified the annotations that were involved in brain or behavioral processes. Those annotations were then categorized by function within each comparison group. We identified 9 functional groups: plasticity (red), DNA/RNA/Transcription (blue), Axon/Dendrite/Synapse (yellow), Ion channel/Receptor (purple), Neuropeptides (green), Immune function (orange), Metabolism (brown), Glia (white), and Other (gray). We saw differences in the relative distribution of GO annotation functional groups across the comparison groups. Neuropeptides were only seen in the *V vs P* group, whereas the *V vs F* group showed a high number of annotations related to Immune Function. The *P vs F* group contained the largest number of annotations related to Plasticity, DNA/RNA/Transcription, and Axon/Dendrite/Synapse.

**Table 2:**
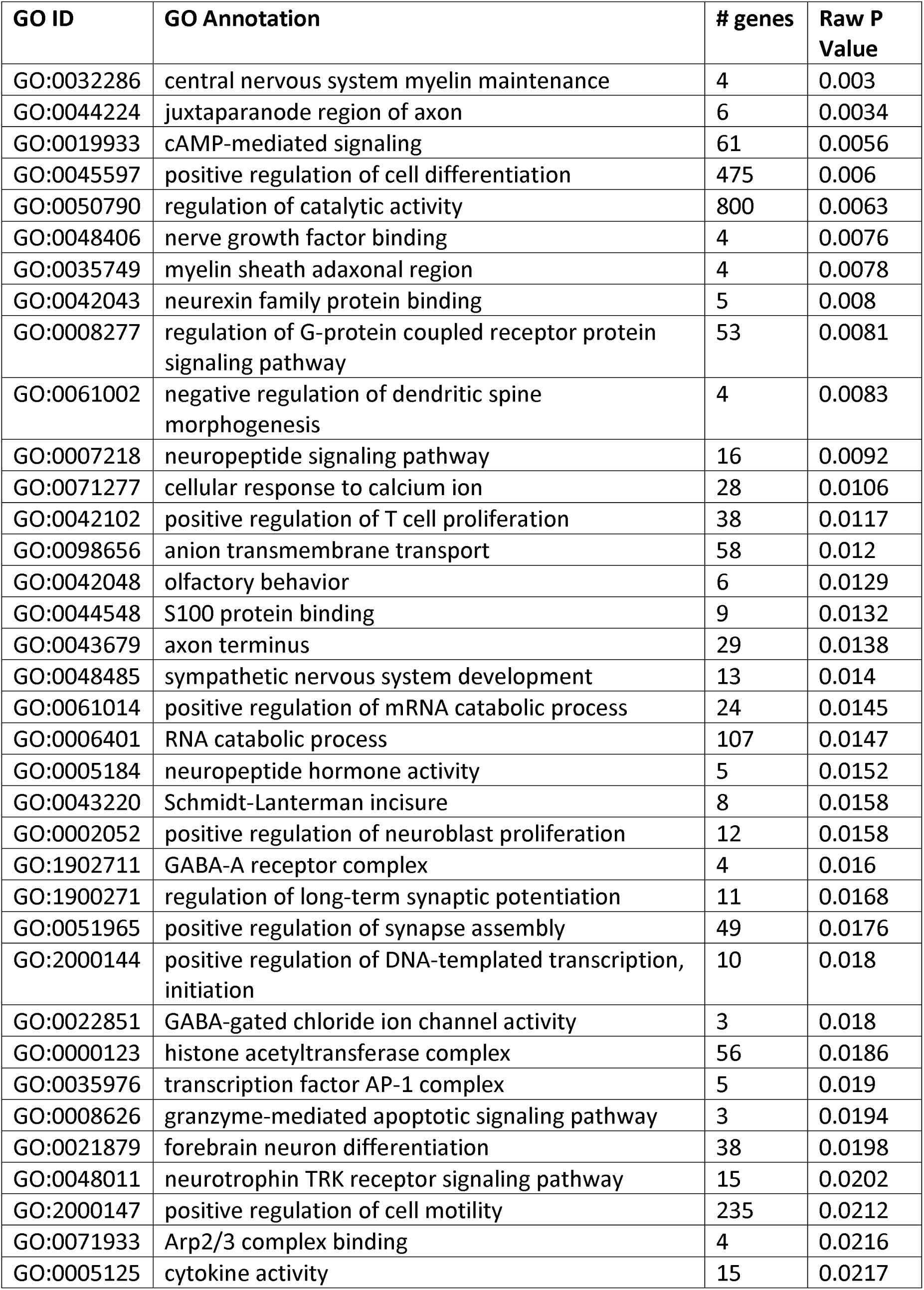
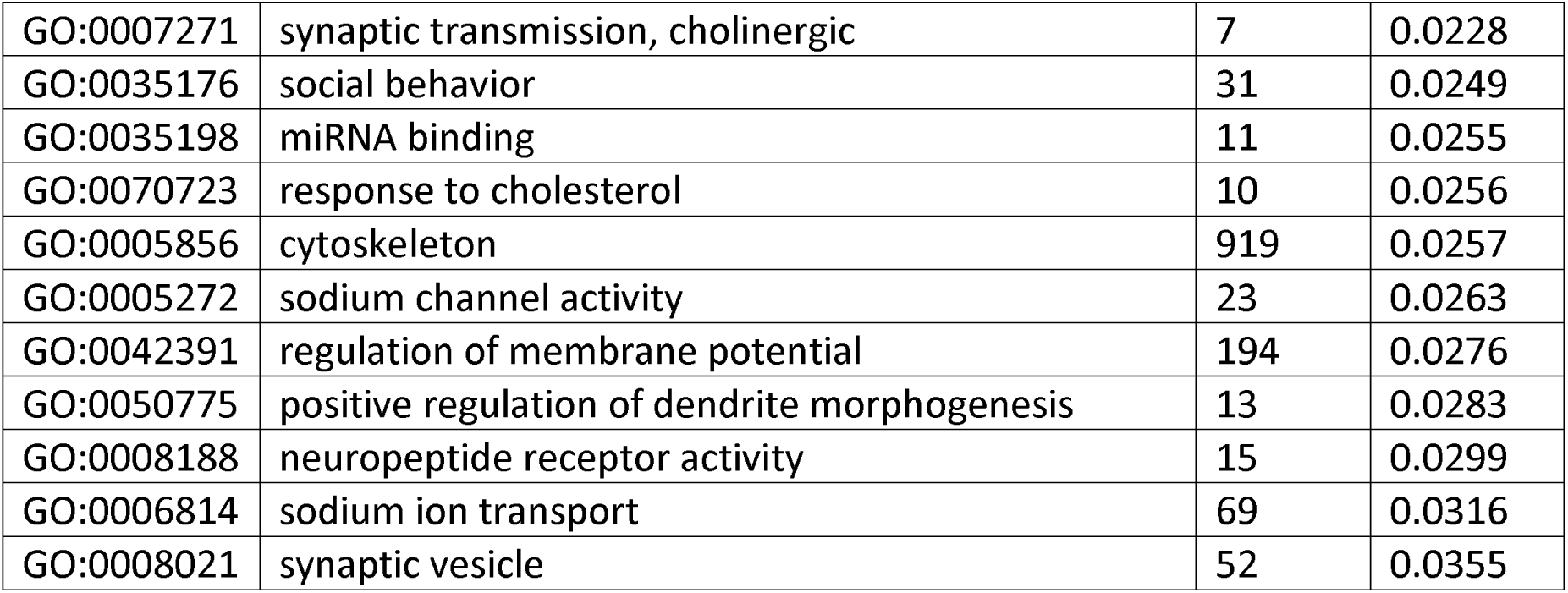
Virgin vs Paired Gene Ontology Annotations

| GO ID | GO Annotation | # genes | Raw P Value |
| --- | --- | --- | --- |
| GO:0032286 | central nervous system myelin maintenance | 4 | 0.003 |
| GO:0044224 | juxtaparanode region of axon | 6 | 0.0034 |
| GO:0019933 | cAMP-mediated signaling | 61 | 0.0056 |
| GO:0045597 | positive regulation of cell differentiation | 475 | 0.006 |
| GO:0050790 | regulation of catalytic activity | 800 | 0.0063 |
| GO:0048406 | nerve growth factor binding | 4 | 0.0076 |
| GO:0035749 | myelin sheath adaxonal region | 4 | 0.0078 |
| GO:0042043 | neurexin family protein binding | 5 | 0.008 |
| GO:0008277 | regulation of G-protein coupled receptor protein signaling pathway | 53 | 0.0081 |
| GO:0061002 | negative regulation of dendritic spine morphogenesis | 4 | 0.0083 |
| GO:0007218 | neuropeptide signaling pathway | 16 | 0.0092 |
| GO:0071277 | cellular response to calcium ion | 28 | 0.0106 |
| GO:0042102 | positive regulation of T cell proliferation | 38 | 0.0117 |
| GO:0098656 | anion transmembrane transport | 58 | 0.012 |
| GO:0042048 | olfactory behavior | 6 | 0.0129 |
| GO:0044548 | S100 protein binding | 9 | 0.0132 |
| GO:0043679 | axon terminus | 29 | 0.0138 |
| GO:0048485 | sympathetic nervous system development | 13 | 0.014 |
| GO:0061014 | positive regulation of mRNA catabolic process | 24 | 0.0145 |
| GO:0006401 | RNA catabolic process | 107 | 0.0147 |
| GO:0005184 | neuropeptide hormone activity | 5 | 0.0152 |
| GO:0043220 | Schmidt-Lanterman incisure | 8 | 0.0158 |
| GO:0002052 | positive regulation of neuroblast proliferation | 12 | 0.0158 |
| GO:1902711 | GABA-A receptor complex | 4 | 0.016 |
| GO:1900271 | regulation of long-term synaptic potentiation | 11 | 0.0168 |
| GO:0051965 | positive regulation of synapse assembly | 49 | 0.0176 |
| GO:2000144 | positive regulation of DNA-templated transcription, initiation | 10 | 0.018 |
| GO:0022851 | GABA-gated chloride ion channel activity | 3 | 0.018 |
| GO:0000123 | histone acetyltransferase complex | 56 | 0.0186 |
| GO:0035976 | transcription factor AP-1 complex | 5 | 0.019 |
| GO:0008626 | granzyme-mediated apoptotic signaling pathway | 3 | 0.0194 |
| GO:0021879 | forebrain neuron differentiation | 38 | 0.0198 |
| GO:0048011 | neurotrophin TRK receptor signaling pathway | 15 | 0.0202 |
| GO:2000147 | positive regulation of cell motility | 235 | 0.0212 |
| GO:0071933 | Arp2/3 complex binding | 4 | 0.0216 |
| GO:0005125 | cytokine activity | 15 | 0.0217 |
| GO:0007271 | synaptic transmission, cholinergic | 7 | 0.0228 |
| GO:0035176 | social behavior | 31 | 0.0249 |
| GO:0035198 | miRNA binding | 11 | 0.0255 |
| GO:0070723 | response to cholesterol | 10 | 0.0256 |
| GO:0005856 | cytoskeleton | 919 | 0.0257 |
| GO:0005272 | sodium channel activity | 23 | 0.0263 |
| GO:0042391 | regulation of membrane potential | 194 | 0.0276 |
| GO:0050775 | positive regulation of dendrite morphogenesis | 13 | 0.0283 |
| GO:0008188 | neuropeptide receptor activity | 15 | 0.0299 |
| GO:0006814 | sodium ion transport | 69 | 0.0316 |
| GO:0008021 | synaptic vesicle | 52 | 0.0355 |

**Table 3:**
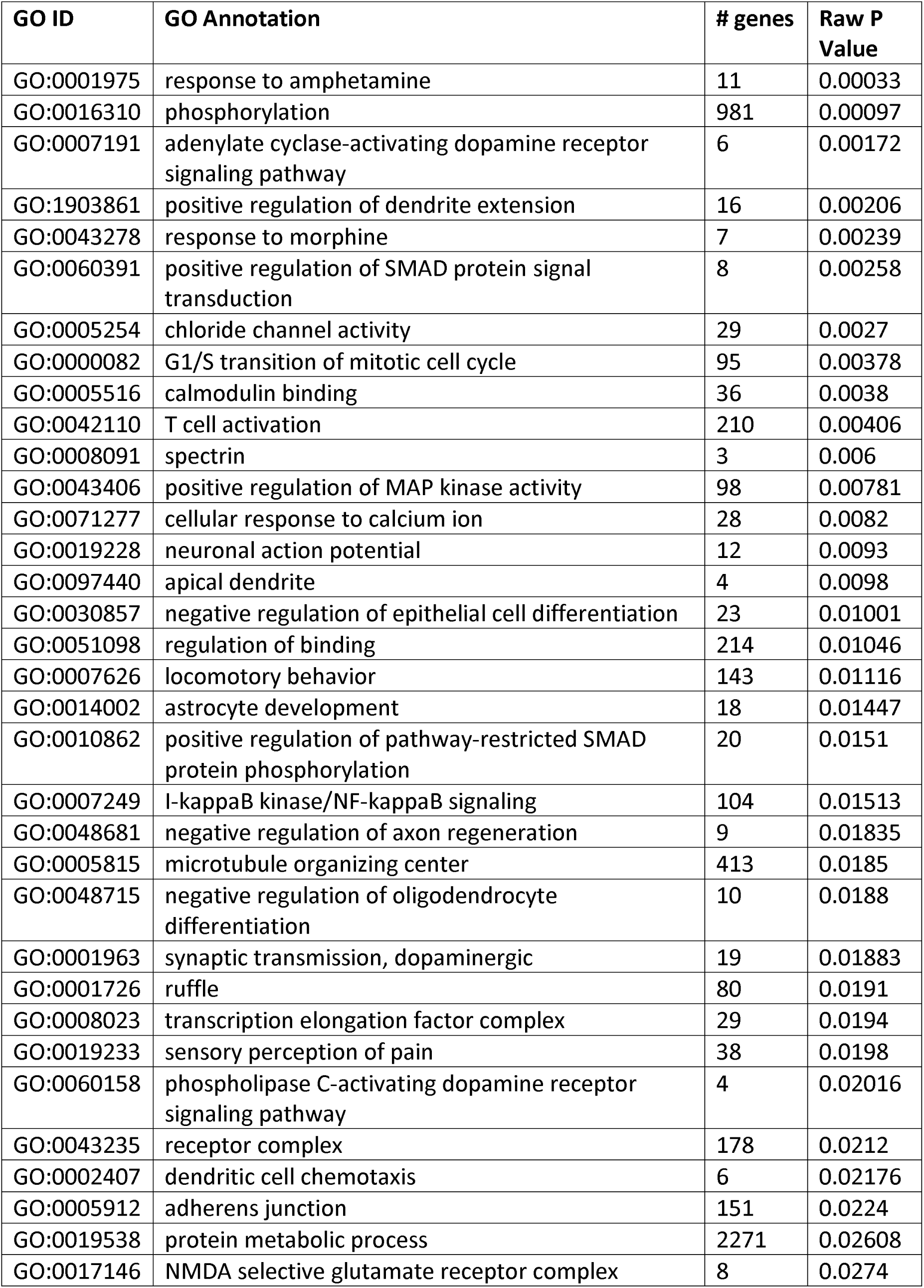
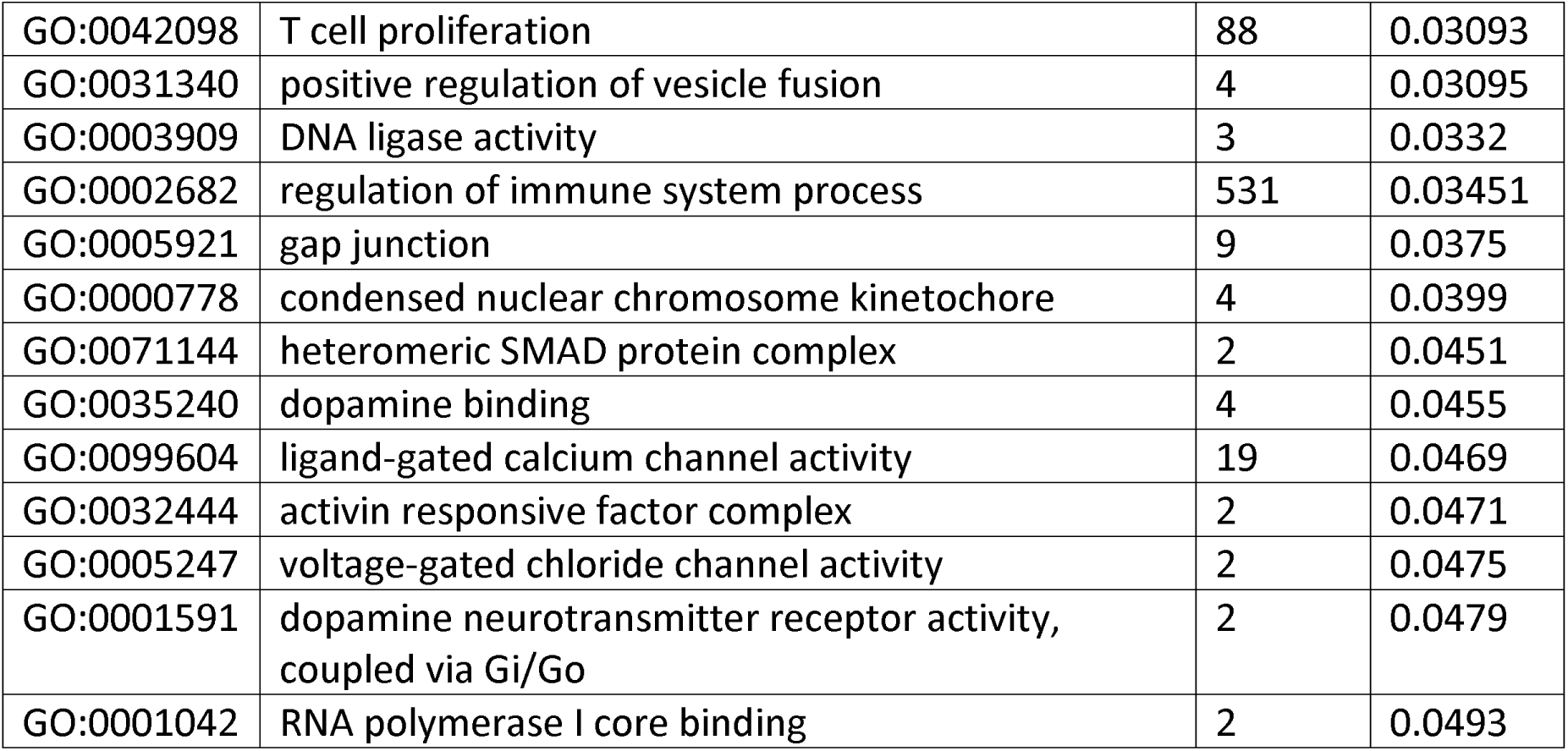
Virgin vs Father Gene Ontology Annotations

**Table 4:** Paired vs Father Gene Ontology Annotations

| GO ID | GO Annotation | # genes | Raw P Value |
| --- | --- | --- | --- |
| GO:0005955 | calcineurin complex | 4 | 0.00104 |
| GO:0050840 | extracellular matrix binding | 33 | 0.00172 |
| GO:0046959 | habituation | 4 | 0.0031 |
| GO:0007626 | locomotory behavior | 144 | 0.0032 |
| GO:2001223 | negative regulation of neuron migration | 7 | 0.004 |
| GO:0046330 | positive regulation of JNK cascade | 64 | 0.0043 |
| GO:0060079 | excitatory postsynaptic potential | 36 | 0.0043 |
| GO:0060391 | positive regulation of SMAD protein signal transduction | 8 | 0.0047 |
| GO:0015116 | sulfate transmembrane transporter activity | 3 | 0.00473 |
| GO:0070723 | response to cholesterol | 10 | 0.0048 |
| GO:0001696 | gastric acid secretion | 6 | 0.005 |
| GO:0051281 | positive regulation of release of sequestered calcium ion into cytosol | 20 | 0.005 |
| GO:0060395 | SMAD protein signal transduction | 38 | 0.0051 |
| GO:0042755 | eating behavior | 10 | 0.0057 |
| GO:0033192 | calmodulin-dependent protein phosphatase activity | 4 | 0.00583 |
| GO:0000403 | Y-form DNA binding | 4 | 0.00644 |
| GO:0010001 | glial cell differentiation | 118 | 0.0065 |
| GO:0048407 | platelet-derived growth factor binding | 11 | 0.0071 |
| GO:0017134 | fibroblast growth factor binding | 11 | 0.00772 |
| GO:0016575 | histone deacetylation | 32 | 0.008 |
| GO:0045893 | positive regulation of transcription, DNA-templated | 783 | 0.009 |
| GO:0005516 | calmodulin binding | 36 | 0.01032 |
| GO:0007616 | long-term memory | 18 | 0.0104 |
| GO:0005882 | intermediate filament | 30 | 0.01192 |
| GO:0061014 | positive regulation of mRNA catabolic process | 24 | 0.0122 |
| GO:0060080 | inhibitory postsynaptic potential | 9 | 0.0124 |
| GO:0035418 | protein localization to synapse | 22 | 0.0141 |
| GO:0008009 | chemokine activity | 4 | 0.01592 |
| GO:0007015 | actin filament organization | 169 | 0.0166 |
| GO:0070410 | co-SMAD binding | 8 | 0.01861 |
| GO:0007212 | dopamine receptor signaling pathway | 20 | 0.0198 |
| GO:0001973 | adenosine receptor signaling pathway | 5 | 0.0199 |
| GO:0005102 | signaling receptor binding | 554 | 0.02035 |
| GO:0000978 | RNA polymerase II proximal promoter sequence-specific DNA binding | 232 | 0.02098 |
| GO:0005736 | DNA-directed RNA polymerase I complex | 7 | 0.02166 |
| GO:0004930 | G-protein coupled receptor activity | 110 | 0.02186 |
| GO:0050882 | voluntary musculoskeletal movement | 6 | 0.0224 |
| GO:0000307 | cyclin-dependent protein kinase holoenzyme complex | 27 | 0.02343 |
| GO:0030374 | ligand-dependent nuclear receptor transcription coactivator activity | 24 | 0.02383 |
| GO:0097110 | scaffold protein binding | 35 | 0.02393 |
| GO:0008622 | epsilon DNA polymerase complex | 3 | 0.02522 |
| GO:0005881 | cytoplasmic microtubule | 29 | 0.02579 |
| GO:0000118 | histone deacetylase complex | 28 | 0.02583 |
| GO:0099061 | integral component of postsynaptic density membrane | 2 | 0.02667 |
| GO:0044309 | neuron spine | 40 | 0.02738 |
| GO:0044295 | axonal growth cone | 7 | 0.02809 |
| GO:0071144 | heteromeric SMAD protein complex | 2 | 0.02824 |
| GO:0043197 | dendritic spine | 37 | 0.02853 |
| GO:0043235 | receptor complex | 179 | 0.0291 |
| GO:0015271 | outward rectifier potassium channel activity | 5 | 0.0317 |
| GO:0042805 | actinin binding | 16 | 0.03185 |
| GO:0003700 | DNA binding transcription factor activity | 385 | 0.03189 |
| GO:0019905 | syntaxin binding | 25 | 0.03526 |
| GO:0098831 | presynaptic active zone cytoplasmic component | 2 | 0.03658 |
| GO:0005794 | Golgi apparatus | 562 | 0.04007 |
| GO:0000976 | transcription regulatory region sequence-specific DNA binding | 368 | 0.04263 |
| GO:0017016 | Ras GTPase binding | 161 | 0.04338 |
| GO:0030864 | cortical actin cytoskeleton | 34 | 0.04449 |
| GO:0014069 | postsynaptic density | 77 | 0.04529 |
| GO:0060053 | neurofilament cytoskeleton | 2 | 0.04637 |
| GO:0008076 | voltage-gated potassium channel complex | 41 | 0.04825 |

### 3.2 Kegg Pathways Analysis

In each comparison group we examined GO annotations with the highest fold enrichment (# observed genes/# expected genes) as well as functions related to neuronal activity, plasticity, or active biological processes, then identified individual genes associated with that function. We ran each individual gene through the Kegg Pathways database (Kyoto Encyclopedia of Genes and Genomes; http://www.genome.jp/kegg/) to identify molecular signaling pathways associated with that gene. Not every gene was associated with a molecular signaling pathway. We then identified commonly recurring pathways associated with the individual differentially expressed genes.

In the *V vs P* comparison group, the commonly recurring pathways included: Protein export, Protein processing in the endoplasmic reticulum, Thyroid hormone synthesis, Antigen processing and presentation, Ras signaling, Rap1 signaling, Neuroactive ligand-receptor pathway, Calcium signaling, and Regulation of the actin cytoskeleton. In the *V vs F* comparison group, the commonly recurring pathways included: Protein processing in the endoplasmic reticulum, Regulation of the actin cytoskeleton, Ras signaling, Metabolic pathways, Axon guidance, Protein processing in the endoplasmic reticulum, Thyroid hormone synthesis, and Antigen processing and presentation. In the *P vs F* comparison group, the commonly recurring pathways included: Ras signaling, Rap1 signaling, Neuroactive ligand-receptor pathway, Calcium signaling, MAPK signaling, LTP, Glutamatergic pathways, Dopaminergic pathways, and Regulation of the actin cytoskeleton.

We next identified genes that were associated with multiple Kegg pathways. By excluding genes that were not associated with any Kegg pathways, or were associated with pathways that were not related to brain function, we further narrowed the range of genes of interest to 49 genes. Ultimately, in each comparison group we identified genes with differential expression across social experience and that were linked to biological pathways within the brain (Table 5). We standardized the expression of each gene relative to its expression in virgin males then grouped genes that were associated with nine commonly recurring Kegg pathways and compared the expression of those genes across groups. Since this was an exploratory study, we did not perform statistical tests, and instead used Cohen’s *d* as a measure of effect size (Figure 5; Table 6). We found large effects of differential expression in genes that were associated with long term potentiation and long term depression (LTP and LTD) (d = 1.072), Neurotransmitters (d = 0.911), and Ca^2+^ Signaling (d = 0.877). We found medium effects of differential expression in genes that were associated with Oxytocin Signaling (d = 0.787), Protein Processing in the Endoplasmic Reticulum (d = 0.599), and Ras/Rap1 Signaling (0.578).

**Table 5:**
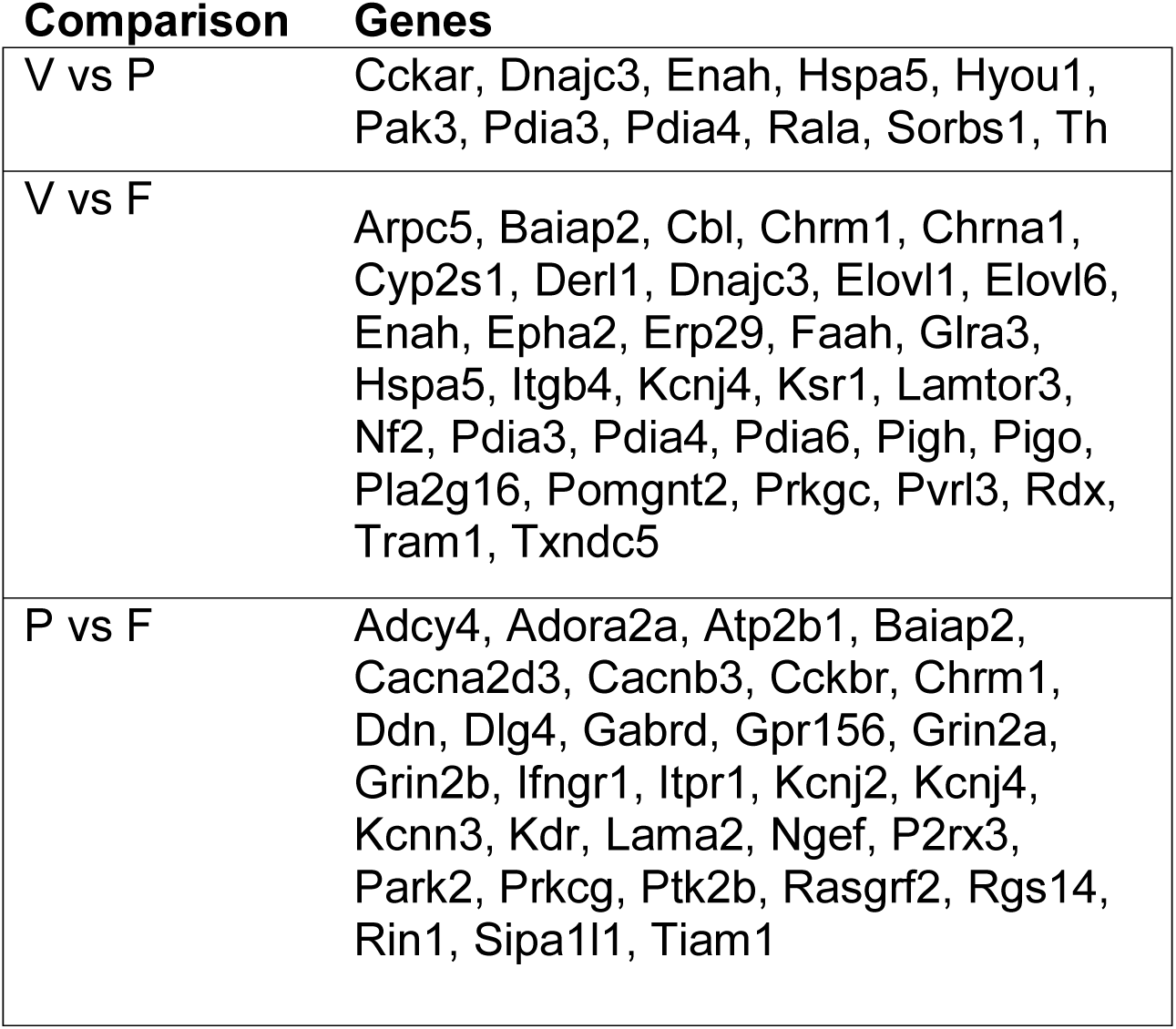
Differentially expressed genes

**Figure 5:**
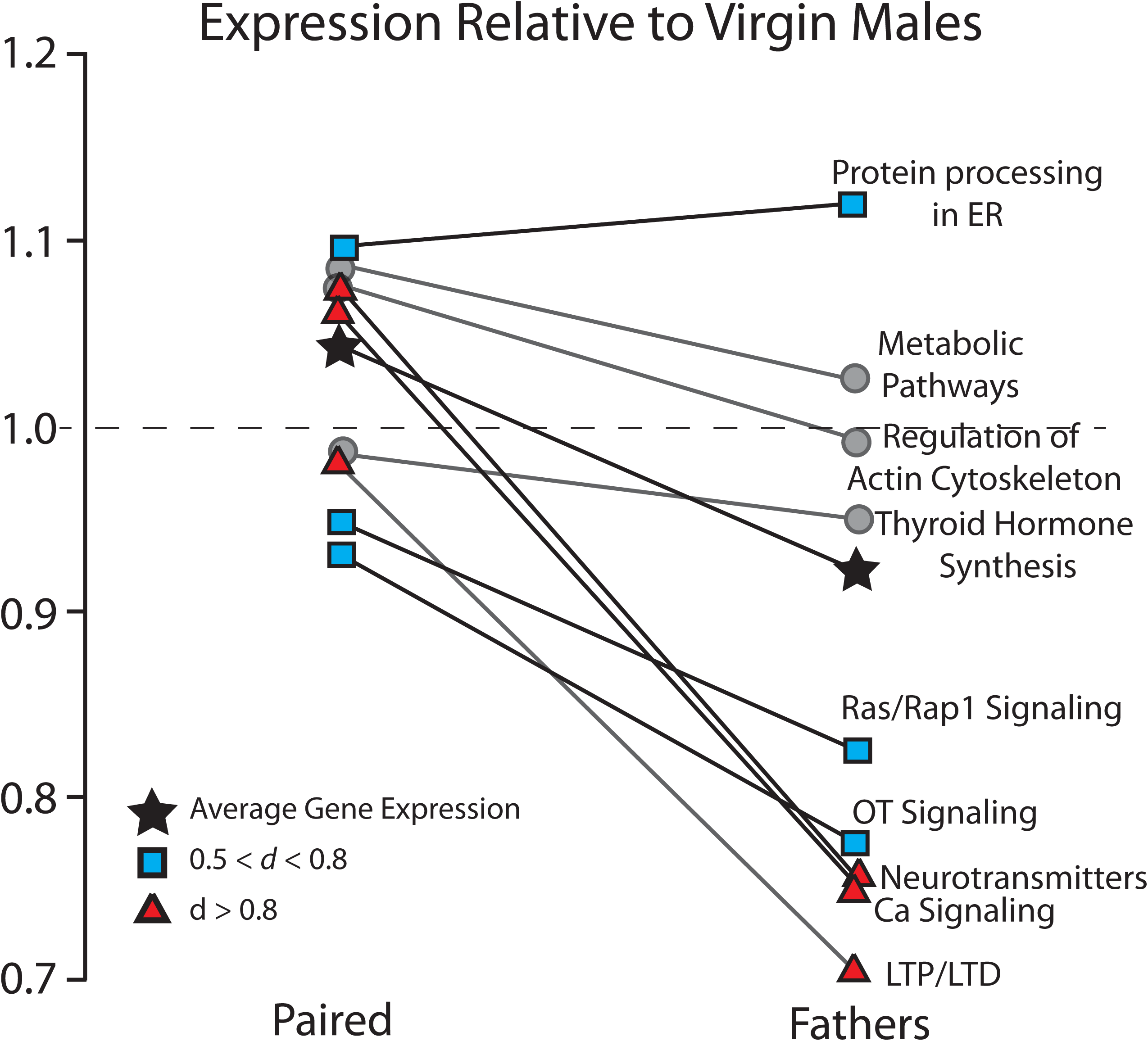
Gene expression in paired males and fathers relative to virgin males. Using the Kegg Pathways analysis, we identified nine pathways of biological or behavioral significance and their associated differentially expressed genes. The mean expression of genes associated with each pathway in fathers was averaged and compared against expression in virgin males. On the whole, gene expression was decreased in fathers relative to both virgins and paired males. Of the nine pathways, only one showed an increase in gene expression in fathers (protein processing in the endoplasmic reticulum), while five showed decreases in gene expression in fathers (Ras/Rap1 signaling, Oxytocin signaling, Neurotransmitters, Calcium signaling, and LTP/LTD). The overall average gene expression is indicated by black stars. Values that exhibited medium effect sizes (0.5 < Cohen’s d < 0.8) are indicated by blue squares and values that exhibited large effect sizes (Cohen’s d > 0.8) are indicated by red triangles).

**Table 6:** Pathways associated with differentially expressed genes

|  |  |
| --- | --- |
| Neurotransmitters | Adcy4, Adora2a, Cacna2d3, Cacnb3, Cckar, Cckbr, Chrm1, Dlg4, Gabrd, Gpr156, Grin2a, Grin2b, Itpr1, Kcnj2, Kcnj4, P2rx3, Prkcg, Rin1 |
| Ca signaling | Adcy4, Adora2a, Cckar, Cckbr, Chrm1, Grin2a, Grin2b, Itpr1, P2rx3, Ptk2b, Rin1 |
| OT signaling | Adcy4, Cacna2d3, Cacnb3, Kcnj2, Kcnj4, Itpr1, Prkcg |
| Regulation of actin cytoskeleton | Arpc5, Baiap2, Chrm1, Enah, Itgb4, Rdx, Tiam1, Enah, Pak3 |
| Ras/Rap1 signaling | Adcy4, Adora2a, Epha2, Grin2a, Grin2b, Kdr, Ksr1, Pak3, Pla2g16, Prkgc, Rala, Rasgrf2, Rgs14, Rin1, Sipa1l1, Tiam1 |
| Protein processing in ER | Derl1, Dnajc3, Erp29, Hspa5, Hyou1, Pdia3, Pdia4, Pdia6, Tram1, Txndc5 |
| Thyroid hormone synthesis | Adcy4, Itpr1, Prkcg, Pdia4, Hspa5 |
| LTP/LTD | Grin2a, Grin2b, Itpr1, Rin1 |

### 3.3 STRING Database Analysis

We used the STRING database to assess the network connectivity between the genes in each comparison group that were identified as having a high degree of differential expression and functional significance. For each analysis, the STRING database constructed a network showing interactions between gene products, as well as the degree of enrichment. The STRING database also performed an enrichment analysis using both Gene Ontology Annotations and Kegg pathways, revealing statistically significant interactions between these gene products.

The 11 differentially expressed genes from the *V vs P* comparison group produced a network with 11 nodes and 11 edges, and a PPI enrichment p-value of 5.86 × 10^−7^ (Figure 6A). Thus, the proteins expressed by these genes have significantly more interactions than would be expected by chance, as defined as a random set of similarly sized proteins selected from the genome. There was one cluster of 7 interacting proteins, and the functions of these gene products were primarily related to functions of the endoplasmic reticulum, as well as the cellular response to stimulation.

**Figure 6:**
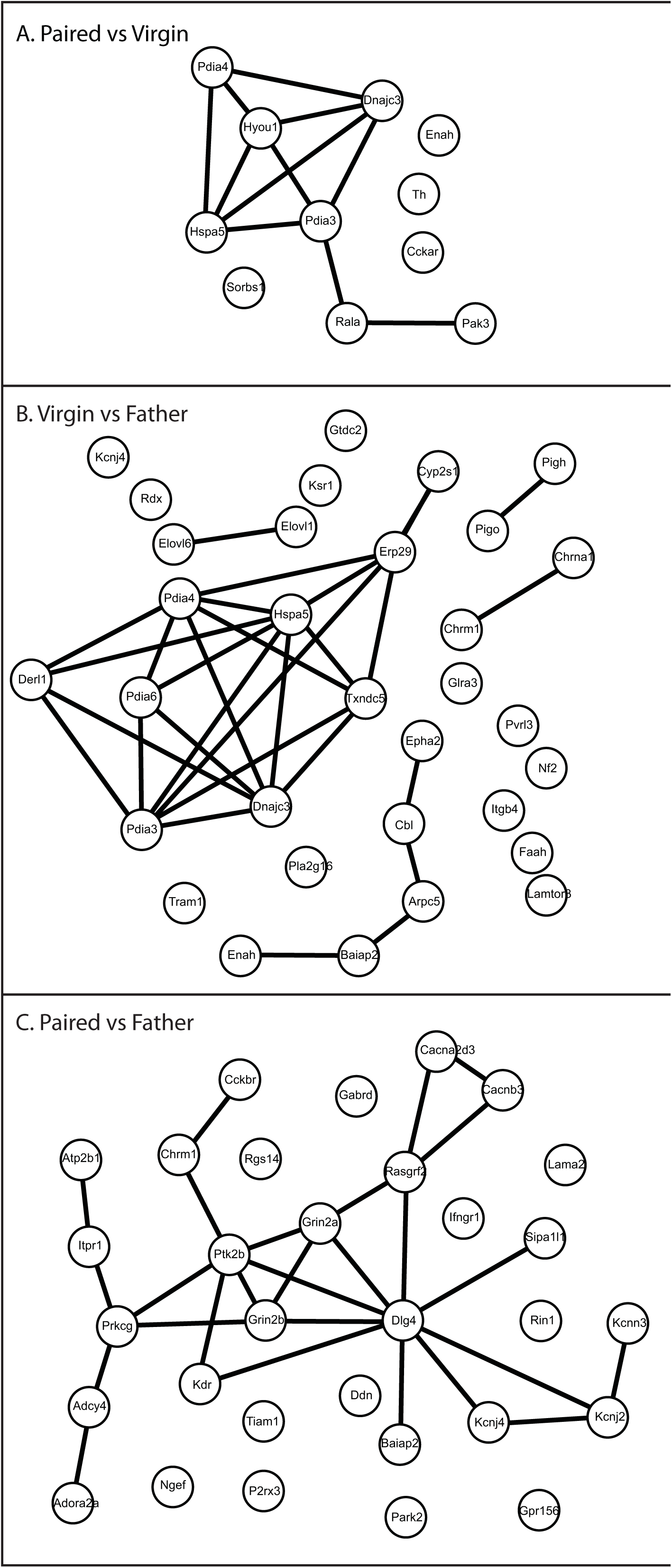
STRING Database analysis of gene product interaction networks. Selected differentially expressed genes were run through the STRING database of gene product interactions, and networks were generated for each comparison. A) Paired vs Virgin network. B) Virgin vs Father. C) Paired vs Father.

The 33 differentially expressed genes from the *V vs F* comparison group produced a network with 32 nodes and 29 edges, and a PPI enrichment p-value of 6.99 × 10^−15^ (Figure 6B), indicating that the proteins expressed by these genes have significantly more interactions than would be expected by chance. These gene products produced one large cluster of 9 interacting proteins, one medium cluster of 5 interacting proteins, and three separate small clusters of 2 interacting proteins. The large cluster was predominantly involved with the function of the endoplasmic reticulum. The medium cluster was involved with process of neural plasticity, including signaling pathways and modification of the actin cytoskeleton. The three small clusters were involved with the elongation of fatty acid chains, the formation of cholinergic receptors, and GPI-anchor synthesis.

The 31 differentially expressed genes from the *P vs F* comparison group produced a network with 31 nodes and 27 edges, and a PPI enrichment p-value of 1.36 × 10^−12^ (Figure 6C), indicating that the proteins expressed by these genes have significantly more interactions than would be expected by chance. These gene products produced one large network consisting of 20 interacting proteins. The genes in this network were involved in a variety of functions, including synaptic plasticity and neural transmission, ion transmembrane transport, the cellular response to stimulus, and the structure of the synapse and dendrite.

### 3.4 NanoString Analysis

A total of 33 genes (30 target genes and 3 housekeeping genes) were selected for quantitative analysis using NanoString. The housekeeping genes (*Gusb*, *Pgk1*, and *Eif4a2*) did not show differential levels of expression across conditions, confirming that these genes can serve as a good baseline in prairie voles. A heat map analysis revealed that 23 of our 30 target genes had lower expression levels in fathers than in either virgins or paired males (Figure 7). Six genes had lower expression levels in virgins, and no gene in any group appeared to show inordinately high levels of expression. A regression analysis revealed similar levels of gene expression across all experimental conditions (Figure 8A).

**Figure 7:**
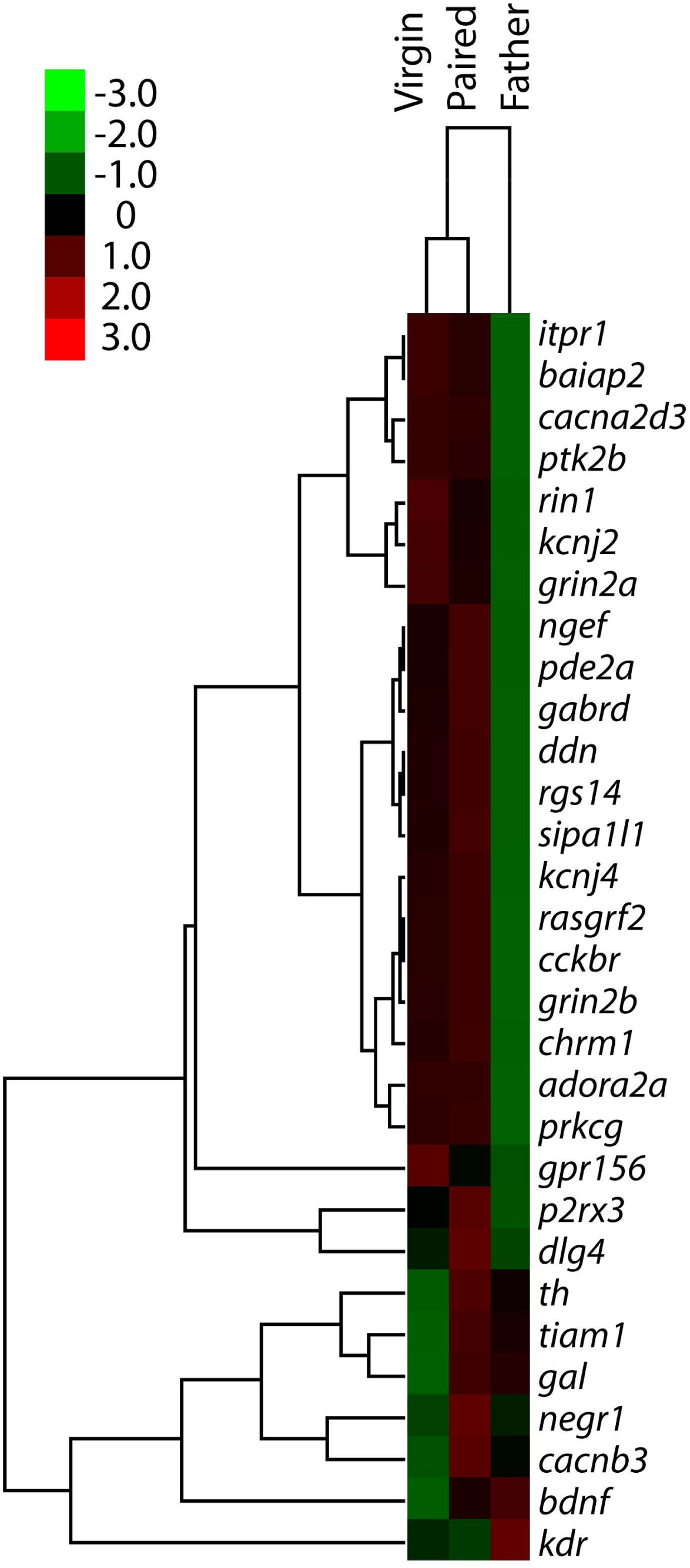
Heat map representing the relative expression of individual genes in virgin males, paired males, and males with fathering experience. Gene enrichment is encoded in the heat map ranging from low (green) to high (red). Genes that show similar expression patterns are clustered together, as indicated by the dendrogram to the left of the heat map.

**Figure 8:**
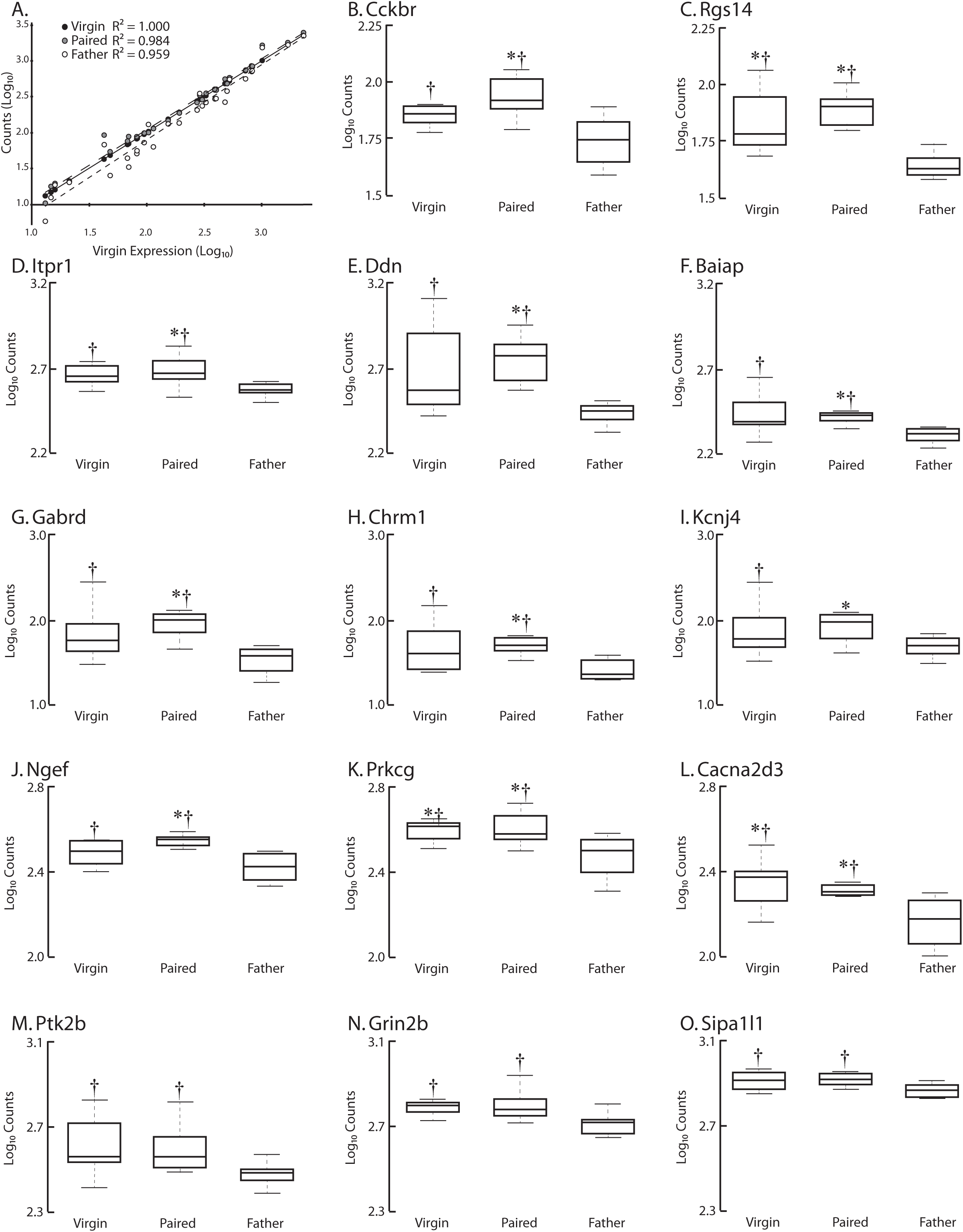
Quantitative analysis of differential gene expression. A) A scatterplot showing B-O) Box and whisker plots showing the expression of genes in virgins, paired males, and fathers. The whiskers represent 1.5x the interquartile range. While we quantified 30 genes, here we show the 14 genes that exhibited significantly different levels of expression across groups or exhibited large effect size. In each gene of these genes, expression in fathers was lower than expression in paired males, and in three cases (Rgs14, Prkcg, and Cacna2d3) expression in fathers was also significantly lower than in virgin males. * - Significantly differs from fathers (p < 0.05). † - Large effect size compared to fathers (Cohen’s d > 0.8).

Expression data for each individual gene was compared across groups using t-tests, which were run and p-values were adjusted for multiple comparisons using nSolver software. Of the 30 target genes, 11 genes showed significant differential expression between comparison groups (p < 0.05; *Cckbr*, *Rgs14*, *Itpr1*, *Ddn*, *Baiap*, *Gabrd*, *Chrm1*, *Kcnj4*, *Ngef*, *Prkcg*, and *Cacna2d3*; Figure 8B-O; Table 7). We also calculated the effect sizes using Cohen’s d, examining differential expression of each gene across groups (Figure 8B-O; Table 8). In the V vs P group, we saw a large effect (defined as 0.8 < d < 1.2) in Tiam1. In the P vs F group, we saw large effects (0.8 < d < 1.2) in *Baiap2*, *Cacna2d3*, *Cckbr*, *Chrm1*, *Ddn*, *Dlg4*, *Gabrd*, *Itpr1*, *Kdr*, *P2rx3*, *Pde2a*, *Ptk2b*, and *Rasgrf2*. In the P vs F group we also saw very large effects (defined as d > 1.2) in *Grin2b*, *Ngef*, *Prkcg*, *Rgs14*, and Sipa1l1. In the V vs F group we saw large effects (0.8 < d < 1.2) in *Adora2a*, *Cacna2d3*, *Cckbr*, *Chrm1*, *Ddn*, *Gabrd*, *Grin2a*, *Grin2b*, *Itpr1*, *Kcnj4*, *Ngef*, *Prkcg*, *Ptk2b*, and Sipa1l1. In the V vs F group we also saw very large effects (d > 1.2) in *Baiap2*, *Rgs14*, and *Rin1*.

**Table 7:** P-values

| Gene Name | F vs P | F vs V | P vs V |
| --- | --- | --- | --- |
| *Adora2a* | 0.1014 | 0.2444 | 0.9891 |
| *Baiap2* | **0.0014** | 0.0776 | 0.7946 |
| *Bdnf* | 0.6273 | 0.2702 | 0.4012 |
| *Cacna2d3* | **0.0460** | **0.0326** | 0.9266 |
| *Cacnb3* | 0.4862 | 0.5735 | 0.2516 |
| *Cckbr* | **0.0120** | 0.0767 | 0.7745 |
| *Chrm1* | **0.0050** | 0.0876 | 0.7353 |
| *Ddn* | **0.0021** | 0.0829 | 0.6383 |
| *Dlg4* | 0.2146 | 0.6877 | 0.2711 |
| *Gabrd* | **0.0018** | 0.0803 | 0.5384 |
| *Gal* | 0.6783 | 0.2519 | 0.1944 |
| *Gpr156* | 0.8590 | 0.7849 | 0.8507 |
| *Grin2a* | 0.2294 | 0.1292 | 0.7381 |
| *Grin2b* | 0.0778 | 0.0877 | 0.7291 |
| *Itpr1* | **0.0500** | 0.0906 | 0.8406 |
| *Kcnj2* | 0.3466 | 0.1956 | 0.7144 |
| *Kcnj4* | **0.0474** | 0.2362 | 0.8533 |
| *Kdr* | 0.2402 | 0.3649 | 0.8613 |
| *Negr1* | 0.5673 | 0.8722 | 0.4501 |
| *Ngef* | **0.0061** | 0.1282 | 0.5444 |
| *P2rx3* | 0.0605 | 0.3671 | 0.2208 |
| *Pde2a* | 0.0677 | 0.2581 | 0.6827 |
| *Prkcg* | **0.0427** | **0.0390** | 0.8554 |
| *Ptk2b* | 0.0729 | 0.0996 | 0.9024 |
| *Rasgrf2* | 0.1027 | 0.1289 | 0.8217 |
| *Rgs14* | **0.0002** | **0.0237** | 0.4867 |
| *Rin1* | 0.1646 | 0.0696 | 0.2535 |
| *Sipa1l1* | 0.0612 | 0.1155 | 0.5573 |
| *Th* | 0.4437 | 0.4343 | 0.1845 |
| *Tiam1* | 0.7299 | 0.2976 | 0.2510 |

**Bold** indicates significant values

**Table 8:** Cohen’s d values

| Gene Name | F vs P | F vs V | P vs V |
| --- | --- | --- | --- |
| *Adora2a* | 0.5380 | **0.9243** | 0.3863 |
| *Baiap2* | **0.9039** | **1.2258** | 0.3220 |
| *Bdnf* | 0.2066 | 0.6260 | 0.4194 |
| *Cacna2d3* | **1.0766** | **1.1004** | 0.0239 |
| *Cacnb3* | 0.4467 | 0.3052 | 0.7520 |
| *Cckbr* | **0.9881** | **0.9884** | 0.0003 |
| *Chrm1* | **0.9371** | **1.0050** | 0.0679 |
| *Ddn* | **1.0688** | **1.0685** | 0.0003 |
| *Dlg4* | **0.8759** | 0.1721 | 0.7037 |
| *Gabrd* | **0.9480** | **0.9667** | 0.0187 |
| *Gal* | 0.2902 | 0.4687 | 0.7589 |
| *Gpr156* | 0.0103 | 0.2844 | 0.2947 |
| *Grin2a* | 0.6531 | **0.8749** | 0.2218 |
| *Grin2b* | **1.2351** | **1.0075** | 0.2276 |
| *Itpr1* | **0.8004** | **1.0156** | 0.2152 |
| *Kcnj2* | 0.5576 | 0.7754 | 0.2178 |
| *Kcnj4* | 0.6183 | **0.8591** | 0.2407 |
| *Kdr* | **0.8634** | 0.6950 | 0.1683 |
| *Negr1* | 0.3141 | **0.1085** | 0.4227 |
| *Ngef* | **1.1987** | **0.9933** | 0.2054 |
| *P2rx3* | **1.1447** | 0.4699 | 0.6748 |
| *Pde2a* | **0.9381** | 0.7403 | 0.1978 |
| *Prkcg* | **1.2933** | **1.1543** | 0.1389 |
| *Ptk2b* | **0.8524** | **0.9836** | 0.1313 |
| *Rasgrf2* | **0.9509** | 0.7855 | 0.1654 |
| *Rgs14* | **1.5018** | **1.2370** | 0.2648 |
| *Rin1* | 0.7439 | **1.4011** | 0.6572 |
| *Sipa1l1* | **1.2207** | **0.9080** | 0.3127 |
| *Th* | 0.3234 | 0.4063 | 0.7297 |
| *Tiam1* | 0.3216 | 0.5269 | **0.8484** |

**Bold** indicates large effect ( $0.8 < d < 1.2$ )
***Bold italics*** indicates very large effect ( $d > 1.2$ )

## 4. Discussion

The transition to fatherhood is associated with a variety of environmental and behavioral changes. In this experiment we sought to identify alterations in central nervous system gene expression that are associated with fathering experience. To our knowledge, this is the first time that RNA sequencing has been performed on prairie vole brain tissue. As such, we faced several technical challenges over the course of this study. For instance, while the prairie vole genome has been sequenced (McGraw et al., 2010; McGraw et al., 2011), its annotation is incomplete, leaving us to rely on the annotated mouse genome (*Mus musculus*) for many of our analyses. In addition, it is important to consider the consequences associated with working in an outbred rodent like prairie voles. The individual differences associated with an outbred population may have masked additional target genes associated with the onset of paternity. Regardless, we still observed significantly altered expression on both the individual gene and system level. These results suggest that paternity engages similar physiological mechanisms across males despite genetic diversity.

Biparental care is rare in mammals, but prairie voles are not the only rodents who exhibit this behavior. The males of several species of *Peromyscus*, including *Peromyscus californicus* and *Peromyscus polionotus*, exhibit paternal care, while other species, including *Peromyscus maniculatus*, do not. This behavioral distinction allowed Bendesky and colleagues to investigate genetic differences between *P. polionotus* and *P. maniculatus* that are linked to parenting behavior (Bendesky et al., 2017). In a series of experiments, they identified several quantitative trait loci that were linked to specific behaviors of interest, including nest building. Further analysis revealed that the gene for arginine vasopression (AVP) was directly related to nest building, and when AVP was administered intracerebroventricularly there was a significant decrease in the quality of nest building (Bendesky et al., 2017). Unlike the study by Bendesky and colleagues, we did not find changes implicating AVP. However, there are several differences between the two experiments. In this study, we specifically examined gene expression within one hypothalamic nucleus, the MPOA. Our study was in a different species and used males that had very specific social experiences: virgin males, pair bonded males, and males with fathering experience.

We targeted the MPOA specifically because it has long been understood to play a role in maternal behavior, and is also believed to be involved in paternal behavior. Lesions to the MPOA disrupt parental behavior in both male and female California mice (Lee and Brown, 2002). In California mouse males, testosterone levels within the MPOA vary in response to parental status (Trainor et al., 2003). Likewise, male California mice with fathering experience show increased Fos-like immunoreactivity in the MPOA following pup exposure (de Jong et al., 2009). To our knowledge, this is the first time that gene expression in the MPOA has been analyzed in the context of fathering behavior.

RNA sequencing is a powerful technique that allows us to identify alterations in gene expression that are associated with behavioral and other phenotypic changes (Wang et al., 2009). The greatest challenge with this technique, however, is the large amount of data it produces. There is no one agreed upon analysis that most effectively identifies specific genes of interest (Conesa et al., 2016; Zhang et al., 2014). Thus, in this study we used several techniques to reveal novel gene targets to further our understanding of paternal behavior. We believe that this is a strength rather than a weakness. All of the target genes identified in this experiment are associated with the experimental differences in social experience. The ultimate goal of this experiment was to increase our understanding of the alterations that occur within the MPOA following exposure to different social contexts in male prairie voles. As such, we have identified a set of genes and their associated pathways that we can use to further explore male parenting behavior.

Our quantitative assessment of gene expression revealed an overall decrease in the expression of many genes in fathers relative to both virgins and pair-bonded males. The specific genes of interest that we identified were involved in a range of physiological processes, including metabolism, stress responsiveness, and plasticity. However, most of the genes that showed significant differential expression, and specifically decreased expression, were associated with synaptic transmission and dendritic spine motility (Table 9). For example, several genes involved in the production and maintenance of receptors (including *Cckbr*, *Chrm1*, *Gabrd*, *Grin2b*, and *Itpr1*) and ion channels (including *Cacna2d3*, *Kcnj4*, and *P2rx3*) were significantly downregulated. These results suggest that GABA, glutamate, and cholinergic systems are all affected by fathering experience, as are calcium and potassium channels. Other genes that exhibited significant downregulation in fathers were involved with the actin cytoskeleton, dendritic spine motility, and other components of the physical plasticity of dendrites. We emphasize that this is not an exhaustive list of differentially expressed genes, however, these results suggest that synaptic plasticity may be diminished in the MPOA of male prairie voles with fathering experience.

**Table 9:** Genes of interest

| Gene ID | Gene Name | Function | GO Annotations |
| --- | --- | --- | --- |
| Baiap2 | Brain-specific angiogenesis inhibitor 1-Associated protein 2 | Insulin receptor tyrosine kinase substrate | Signaling, Regulation of Biological Quality, Membrane Part, Dendrite, Dendritic Spine, Synapse |
| Cacna2d3 | Calcium voltage gated channel, auxiliary subunit alpha 2 delta 3 | Voltage gated calcium channel | Ion Channel Activity |
| Cckbr | Cholecystokinin B receptor | Multipass transmembrane receptor protein | Signaling, Regulation of Biological Quality |
| Chrm1 | Cholinergic receptor, muscarinic 1 | Muscarinic receptor | Signaling, Regulation of Biological Quality, Membrane Part, Dendrite, Synapse |
| Ddn | Dendrin | plasma membrane surrounding dendritic spine | Membrane Part |
| Gabrd | GABA A receptor, subunit delta | GABA receptor | Signaling, Dendrite, Synapse, Ion Channel Activity |
| Grin2b | Glutamate receptor, ionotropic, NMDA 2b | NMDA receptor | Signaling, Regulation of Biological Quality, Membrane Part, Dendrite, Dendritic Spine, Synapse, Ion Channel Activity |
| Itpr1 | inositol 1,4,5-triphosphate receptor 1 | Calcium channel | Signaling, Regulation of Biological Quality, Membrane Part, Dendrite, Synapse, Ion Channel Activity |
| Kcnj4 | potassium voltage gated channel subfamily J member 4 | potassium channels - ion homeostasis | Membrane Part, Dendrite, Synapse, Ion Channel Activity |
| Ngef | Neuronal guanine nucleotide exchange factor | dendritic spine morphogenesis | Signaling, Membrane Part |
| P2rx3 | purinergic receptor p2x, ligand gated ion channel | ATP receptor | Signaling, Regulation of Biological Quality, Membrane Part, Dendrite, Dendritic Spine, Synapse, Ion Channel Activity |
| Pde2a | Phosphodiesterase 2a | 2nd messenger signaling/dendritic spines | Signaling, Regulation of Biological Quality, Membrane Part, Dendrite |
| Prkcg | protein kinase c gamma | signaling protein | Signaling, Membrane Part, Dendrite, Synapse |
| Ptk2b | protein tyrosine kinase 2 beta | ion channel regulation; MapK signaling | Signaling, Regulation of Biological Quality, Membrane Part, Dendrite, Dendritic Spine, Synapse, Ion Channel Activity |
| Rgs14 | regulator of g-protein signaling 14 | scaffold protein | Signaling, Regulation of Biological Quality, Membrane Part, Dendrite, Dendritic Spine, Synapse |
| Rin1 | Ras and Rab interactor 1 | Ras effector | Signaling, Regulation of Biological Quality, Membrane Part, Dendrite |
| Sipa1l1 | signal induced proliferation<br>associate 1 like 1 | Ras effector | Signaling, Regulation of Biological Quality,<br>Membrane Part, Dendrite, Dendritic Spine,<br>Synapse |

We were surprised by the lack of differential expression of oxytocin and vasopressin related genes, however this finding is not unique within the literature. In a series of experiments, Kenkel and colleagues examined the neuroendocrine correlates of pup exposure in male prairie voles that were virgins or had fathering experience (Kenkel et al., 2012; Kenkel et al., 2014). They saw changes in OT immunoreactivity in PVN/BNST, but there were no changes to OT/AVP in the MPOA. Another study examined OT immunoreactive cells in male prairie voles that were virgins, had established pair bonds, or had fathering experience (Wang et al., 2015). They saw an increase in the number of OT immunoreactive cells in the MPOA of paired males and fathers compared to virgin males, but there was a greater increase of OT-ir cells in the PVN of fathers compared to paired and virgin males. It is likely that examination of gene expression in the PVN would show alterations in OT gene expression. In future studies we hope to examine these and other brain regions.

Fatherhood also seems to be associated with structural alterations in neural plasticity, as measured by changes in the number and density of dendritic spines. Mice with fathering experience show increased survival of newborn neurons and increased dendritic spine density within the hippocampus (Glasper et al., 2016; Hyer et al., 2016). Male marmosets show an increase in dendritic spine density in the prefrontal cortex after fathering experience (Kozorovitskiy et al., 2006). However, other studies have shown reductions in the survival of adult-generated neurons in the amygdala and hippocampus (Glasper et al., 2011; Lieberwirth et al., 2013). It seems clear that the effects of fatherhood vary across brain regions, but we do not yet know what is causing these changes in neural plasticity.

The lower gene expression related to dendritic spines, shown to be associated with fatherhood in the present study, is evocative of similar changes seen in a recent study of the MPOA of mother rats (Parent et al., 2017). *Rem2*, a gene associated with reduction of dendritic branching but increases in spine density (Ghiretti et al., 2014; Ghiretti and Paradis, 2011) was increased in the MPOA of high licking/grooming rats, but only in lactating mothers (not in virgins). This increase was accompanied by decreased dendritic complexity. *Rem2* is involved with GTPase activity and GTP binding. While we did not see alterations in *Rem2* expression in this study, we found differential expression of several genes that are involved in Ras and Rap1 signaling. Both Ras and Rap1 are GTPases that play an important role in the structural and functional plasticity of cells (Cahill et al., 2016; Zhu et al., 2002).

The down-regulation of genes associated with dendritic complexity in the present study, as well as the study by Parent and colleagues, is similar to what one would expect in an animal that had experienced high amounts of stress. It is well established that stress, mediated by corticotropin-releasing hormone, results in a loss of dendritic spines (Chen et al., 2008; Chen et al., 2013; Leuner and Shors, 2013; Liao et al., 2014; Radley et al., 2006). Interestingly, rat mothers show a decrease in the number and density of dendritic spines in the amygdala and stria terminalis four days after birth (Matsuo et al., 2017), and an increase in dendritic spine density in the hippocampus during the postpartum period (Kinsley et al., 2006). This suggests that alterations in dendritic spine density in mothers are both transient and region specific. More studies must be done to determine if the same holds true for vole fathers.

In many species, the transition to fatherhood is associated with a suite of behavioral and hormonal changes, including those indicative of stress. In California mice (*Peromyscus californicus*), fathers exhibit attenuated anxiety-like behavior approximately two weeks after pups are born (Glasper et al., 2016; Hyer et al., 2016). Human males show a peak in cortisol levels during the transition to fatherhood (Storey et al., 2000), but this is highly variable across studies (Gordon et al., 2010). Prairie vole fathers show increased anxiety-like behavior, and chronic pup exposure results in an increase in basal CORT levels (Lieberwirth et al., 2013). In an open field test, fathers spent more time in corner squares, and in an elevated plus maze, fathers spent less time in open arms. In forced swim tests, fatherhood decreased the latency to immobility, and increased the number and duration of immobility bouts (Lieberwirth et al., 2013). Additionally, males with fathering experience showed fewer BrdU-labeled cells in the amygdala, hippocampus, and ventromedial hypothalamus than virgin males (Lieberwirth et al., 2013). In the long term, fatherhood may be beneficial for male health, but the transition to fatherhood is a tremendously stressful period (Bartlett, 2004).

In male voles with fathering experience we also see the upregulation of genes related to protein processing in the endoplasmic reticulum. The endoplasmic reticulum is instrumental in managing the protein folding process, including disposing of misfolded proteins (Zhang and Kaufman, 2008). Homeostatic imbalances, including stress, can alter the functioning of the endoplasmic reticulum, leading to the initiation of the unfolded-protein response, which can in turn lead to apoptosis (Banhegyi et al., 2007; Mandl et al., 2009). This may be one mechanism by which physiological stress can result in homeostatic perturbations (Walter and Ron, 2011; Zhang and Kaufman, 2006), including some of the changes that are evident in vole fathers, such as weight loss (Campbell et al., 2009; Kenkel et al., 2014).

While many of the changes we saw in gene expression may be partially attributable to stress, there are likely many other additional factors at play. Fathers in many species show systematic endocrine changes (Saltzman and Ziegler, 2014). Environmental factors, including changes in the types and amount of sensory stimulation, or the amount of parental care they received, may play a role as well (Braun and Champagne, 2014; Champagne, 2016). Much more work must be done to tease apart these many factors.

In this study we saw the most varied and interesting differences between the paired males and males with fathering experience. This was surprising, as we expected that the greatest differences would be between the virgin males and fathers. However, examination of the quantitative results begins to clarify these findings (Figs 5 and 8). The expression of genes of interest is slightly elevated in paired animals relative to virgins, but the expression in fathers is decreased relative to virgins. Thus, while the expression levels of some genes do not significantly differ between virgins and paired males, and virgins and fathers, we found significant differences between paired males and fathers. This may suggest that the experience of fathering is functionally distinct from any other type of social interactions that these animals have encountered.

### 4.1 Conclusions

The purpose of this study was to explore how gene expression changed across the transition to fatherhood, and to identify novel targets to allow for deeper investigation of male parenting behavior. The use of RNA sequencing confirmed that there are differences in gene expression between voles that had different social experiences, including virgin males, males that had formed a pair bond with a female, and males with parenting experience. The genes identified in this study suggest novel processes that are related to paternal behavior and offer new targets for the further exploration of fathering behavior.

## Acknowledgments

The sequencing was carried out at the DNA Technologies and Expression Analysis Cores at the UC Davis Genome Center, supported by NIH Shared Instrumentation Grant 1S10OD010786-01, and was made possible by a pilot grant from the UC Davis Genome Center. Thanks to Chris Harshaw for helpful analytical suggestions. Thanks to Cindy Clayton and Rhonda Oates-O’Brien for husbandry and veterinary care of the prairie vole colony. And special thanks to the anonymous reviewers for their helpful and constructive comments on this manuscript.

